# Daily vocalization patterns of the Saipan Reed Warbler (*Acrocephalus hiwae*)

**DOI:** 10.1101/2024.02.02.578722

**Authors:** Willson Gaul, Jie Lin, Ellie Roark

**Affiliations:** Department of Science, Mathematics, Health, and Athletics, Northern Marianas College, Saipan, Commonwealth of the Northern Mariana Islands, USA; Division of Fish and Wildlife, Commonwealth of the Northern Mariana Islands, USA

**Keywords:** automated recording unit, diel vocalization, endangered species, tropical bird song, Mariana Islands

## Abstract

We investigated how detectability and vocalization patterns of Saipan Reed Warblers (*Acrocephalus hiwae*) varied by time of day. We used long-duration sound recordings from eleven locations occupied by Saipan Reed Warblers to model the probability of detecting a vocalization in each hour of the day. We found that Saipan Reed Warblers sang during all daylight hours. We did not find evidence of a dawn or evening chorus in this species. These results are useful for determining what time of day surveys of Saipan Reed Warblers should be conducted, which is particularly relevant because Saipan Reed Warblers are protected by local and U.S. Federal laws.

---

The genus *Acrocephalus* is a group of Old World Reed Warblers that includes many island endemic species (Winkler et al. 2020). The Mariana Islands are an archipelago of volcanic and uplifted limestone islands in the western Pacific Ocean (Cloud et al. 1957). At least two of the Mariana Islands have extant populations of Reed Warblers in the genus *Acrocephalus*, while four *Acrocephalus* species have been extirpated from four other islands in the chain (Saitoh et al. 2012). The extant populations on the islands of Saipan and Alamagan are considered by Saitoh et al. (2012) to belong to the same species, the Saipan Reed Warbler *Acrocephalus hiwae*. The local common names for this bird are Saipan Reed Warbler (English), Ga’ ga’ karisu (Chamorro), and Malul ghariisu (Carolinian) (Northern Mariana Islands Administrative Code § 85-30.1-101 2016). The Saipan Reed Warbler populations on Saipan and Alamagan are listed as threatened or endangered under Commonwealth of the Northern Mariana Islands (CNMI) and U.S. law under the name Nightingale Reed Warbler, *Acrocephalus luscinia* (Northern Mariana Islands Administrative Code § 85-30.1-101, 2016; Conservation of endangered 1970).

The Saipan Reed Warbler is a high priority species for conservation and management activities by both local and federal organizations because it is a locally and federally protected species. The current global population of Saipan Reed Warblers is estimated to be between 1,019 and 6,356 individuals, with approximately 75% residing on Saipan, and 25% on Alamagan (Marshall et al. 2021). Surveys of the Saipan population found lower Saipan Reed Warbler density in 2007 than in 1982 (Camp et al. 2009), while the Alamagan population showed no significant difference in density between 2000 and 2010 (Marshall et al. 2021). Local agencies and their partners are currently preparing to translocate Saipan Reed Warblers to other islands in the Mariana archipelago to increase its range-wide resilience to anthropogenic threats (CNMI DFW 2014). Threats to the species include limited distribution, habitat conversion, Brown Tree Snake (*Boiga irregularis*) and other invasive predators, and stochastic events such as typhoons (Marshall et al. 2021).

Surveys to determine Saipan Reed Warbler presence are regularly conducted by government agencies and consultancies to ensure development projects are in compliance with the U.S. Endangered Species Act and local laws. Building or development that results in take of Saipan Reed Warblers may be offset by purchasing credits that fund habitat protection in the Saipan Upland Mitigation Bank, a forested area of approximately 330 hectares on the northern part of Saipan (Northern Mariana Islands Public Law No. 10-84 1997). In addition to surveys for development, the CNMI Division of Fish and Wildlife (DFW) also conducts surveys to assess population status of Saipan Reed Warblers on Saipan and Alamagan. Recovery criteria for downlisting Saipan Reed Warblers include maintaining or increasing population numbers for five consecutive years on both Alamagan and Saipan (USFWS 1998). To delist the species, US Fish and Wildlife Service (USFWS) requires that there be at least 8,000 individuals in the Mariana Islands, distributed in stable or increasing populations on at least five islands (USFWS 1998). Increasing the efficiency and efficacy of monitoring efforts is key to evaluating the recovery of this species, and to assessing the effectiveness of conservation actions.

The USFWS recommends 10 replicate visits to survey for Saipan Reed Warbler occupancy (USFWS 2014). The survey window recommended by USFWS is sunrise to “around 10:00 hours” and the two hours before sunset (USFWS 2014). In practice, 95% of all surveys by the CNMI DFW at proposed development sites since 2005 have been conducted between 0707 h and 1510 h (Supplemental Table S1). On Alamagan and Saipan, in areas not targeted for development, DFW surveys for Saipan Reed Warblers have generally been conducted between dawn and 1100 h, using 5-min stationary point counts (or 8-min stationary point counts on Alamagan) along established transects. This is sometimes followed by a period of playback to increase the probability of detecting a bird when it is present (ER pers. obs, Marshall et al. 2021).

Tropical passerines show a variety of breeding patterns and behaviors that are uncommon in temperate regions, including asynchronous breeding, singing and territory defense by both males and females, and year-round territory defense (Stutchbury and Morton 2023). Tropical birds also differ in their daily singing patterns; some species of tropical birds have strong dawn choruses, but other species do not (Stutchbury and Morton 2023).

Understanding the daily singing patterns of Saipan Reed Warblers is important when planning targeted surveys for this species, because surveying at times when the birds are most vocal will increase the probability of detecting the species if it is present.

Automated recording units (ARUs) are compact audio recorders that can be deployed for extended time periods to increase the temporal and spatial extent of auditory bird surveys without increasing the costs associated with human observer hours in the field (Williams et al. 2018). Passive acoustic monitoring is widely used to study many taxa, including bats (Tuneu-Corral et al. 2020), whales (Baumgartner et al. 2019), invertebrates (Penone et al. 2013), and often as a substitute for point counts to monitor birds (Furnas and Callas 2015, Klingbeil and Willig 2015, Shonfeld and Bayne 2017, Darras et al. 2018).

The CNMI DFW and its partners plan to continue Saipan Reed Warbler surveys on Saipan and Alamagan both for development permitting and population monitoring. The CNMI DFW will also need to monitor any potential future translocated populations to meet recovery goals (CNMI DFW 2014). Using ARUs to monitor long-term resident and translocated bird populations in the remote and difficult to access islands north of Saipan in the Mariana archipelago will increase survey effort while minimizing the costs and logistical constraints associated with in-person surveys.

In this study, we used ARUs to investigate the daily vocalization patterns of Saipan Reed Warblers. This study provides novel information about the vocal behavior of Saipan Reed Warblers, and fills a knowledge gap for designing targeted survey protocols for this species.

## Methods

We analyzed the daily vocalization patterns of Saipan Reed Warblers using data collected on the island of Saipan, in the CNMI. During the years 2021 through 2023, we deployed continuously recording ARUs on 15 known or suspected Saipan Reed Warbler territories in both the wet season (July) and the dry season (January-March). We used random forests to model observed vocalization patterns to understand the daily vocalization behavior for this species. Code and data to produce figures and results are available on Zenodo (Gaul 2024).

### Study area

We deployed ARUs on Saipan, primarily in conservation areas or on public land where human activity is relatively limited. While Saipan Reed Warblers also occur in more densely human-inhabited areas, we prioritized collecting sound recordings in areas that allowed us to minimize incidental recording of human activities, and other anthropogenic sources of noise.

We deployed ARUs at 15 locations with suspected Saipan Reed Warbler occupancy. Occupancy was suspected if a Saipan Reed Warbler had been detected there over the previous calendar year by either DFW staff members or the authors. Locations for ARU deployment were selected opportunistically based on land access availability. Recorders were deployed at one location in August 2021, at 11 locations in July 2022, and at 11 locations between January and March of 2023. Deployments in 2023 included 8 previously sampled locations, and 3 new locations. We discarded 4 locations where Saipan Reed Warblers were never detected on ARU recordings, because we were interested in studying vocalization patterns at locations where we were certain that Saipan Reed Warblers were present.

### Sound recordings

Sound recordings were collected using SWIFT bioacoustic automated recording units (Cornell Lab of Ornithology, Ithaca, NY, USA), which used a built-in PUI Audio brand omni-directional microphone. ARUs recorded at a sampling rate of 48 kHz and saved recordings as uncompressed .WAV files. The microphone gain was set to 35 dB, with a signal to noise ratio of approximately 58 dB reported by the manufacturers.

ARUs were programmed to record continuously for 24 h per day during deployment. They were deployed at each site for a minimum of 48 h (often more) to ensure that they recorded at least one 24 h period with minimal noise disturbance (either from anthropogenic sources or from weather conditions). ARUs were deployed by strapping the units to trees at a height of approximately 1.5 m (Darras et al. 2018). ARUs were always attached to trees that were less than 23 cm in diameter to avoid blocking the microphone from sampling behind the tree.

### Audio data processing

Audio recordings were processed using a desk-based listening method, in which an observer listened to randomly selected one-minute segments from extended-duration recordings (Roark and Gaul 2021). A technician listened through headphones and viewed spectrograms of the recordings in Audacity (Audacity Team 2023) to determine whether a Saipan Reed Warbler vocalization was present or absent. Initially, observers inspected 10 minutes from each hour. After approximately 15 hours of recordings had been processed this way, we subsampled this data and determined that inspecting 4 minutes per hour would provide similar results with significantly less data processing time. All subsequent hours were analyzed with 4 randomly selected minutes per hour. Here, we present results using all minutes that were searched by an observer.

For each selected minute of audio, the observer noted whether a Saipan Reed Warbler was detected vocalizing within the selected minute, or whether there was too much noise to determine whether a Saipan Reed Warbler vocalization was present. If there was too much noise to determine whether a vocalization was present, that minute was not used in subsequent analyses. A subset of minutes were searched for Saipan Reed Warbler sounds by 2 observers independently. The observers’ data were compared, and the reliability of Saipan Reed Warbler detections was assessed using Krippendorf’s alpha, which measures agreement between multiple observers after accounting for agreement by chance (Gamer et al. 2019, Krippendorff 2013).

### Statistical analyses

We modeled the probability of detecting a Saipan Reed Warbler vocalization in each hour of the day using random forests (Breiman 2001). The response variable was a dichotomous variable indicating whether a Saipan Reed Warbler vocalization was detected in the hour. The predictor variables were the time of day (in hours), which was the variable of interest, and 2 covariates accounting for sampling skill and effort: the identity of the observer; and the total number of minutes listened to from that hour. We assessed model predictive performance using two metrics: area under the receiver operating characteristic curve (AUC), which measures the ability of a model to discriminate between two classes (Fielding and Bell 1997); and Brier score, which measures model calibration as the mean squared error between predicted probabilities and binary observations (Brier 1950).

We fit random forests using the ‘randomForest’ package in R version 4.3.1 (Liaw and Wiener 2002, R Core Team 2023). We generated 100 bootstrap samples of the data to estimate bootstrap confidence intervals around model prediction performance metrics and pointwise bootstrap confidence intervals around model predictions (Hastie et al. 2009). For each bootstrap sample of the data, we fitted models using 11-fold cross-validation (CV) (Hastie et al. 2009). In each CV fold, data from one location were withheld as a test set and the random forest model was fitted with data from the remaining 10 locations. Prediction performance was measured using predictions to the holdout data in each fold, and we reported mean prediction performance measured from all holdout folds across all bootstrap datasets.

To test whether time of day had a significant effect on the probability of detecting a Saipan Reed Warbler, we fit a model representing the null hypothesis, excluding time of day, so that the only the predictor variables were the identity of the observer and the number of minutes searched. We then compared AUC and Brier score from the models with and without the time of day as a predictor variable.

To visualize the daily vocalization patterns of Saipan Reed Warblers, we generated model predictions to standardized data. The standardized predictions show the predicted probability of our most experienced observer (ER) detecting a Saipan Reed Warbler vocalization in each hour of the day if she listened to 4 randomly selected minutes of audio recording per hour. We graphed standardized predictions averaged over all the bootstrapped, cross-validated model replicates.

## Results

We detected Saipan Reed Warbler vocalizations on recordings from 11 locations. Our final dataset consisted of 3,743 min of audio recording, randomly sampled from 927 h of recordings at 11 locations. Comparison of data from multiple observers showed that identifications of Saipan Reed Warbler sounds were reliable (Krippendorf’s alpha = 0.735, *n* = 215 min searched by 2 observers).

Saipan Reed Warblers began singing around sunrise, continued singing throughout the day, and stopped singing around sunset (Fig. 1). The earliest detected Saipan Reed Warbler sound was just after midnight at 0016 h (ChST), and the latest detected Saipan Reed Warbler sound was at 1907 h, fifteen minutes after sunset. Ninety-five percent of our detections occurred between 0545 h and 1826 h. Saipan Reed Warbler vocalizations were detected in every hour from 0500 h to 1900 h (Fig. 1).

**Figure 1.**
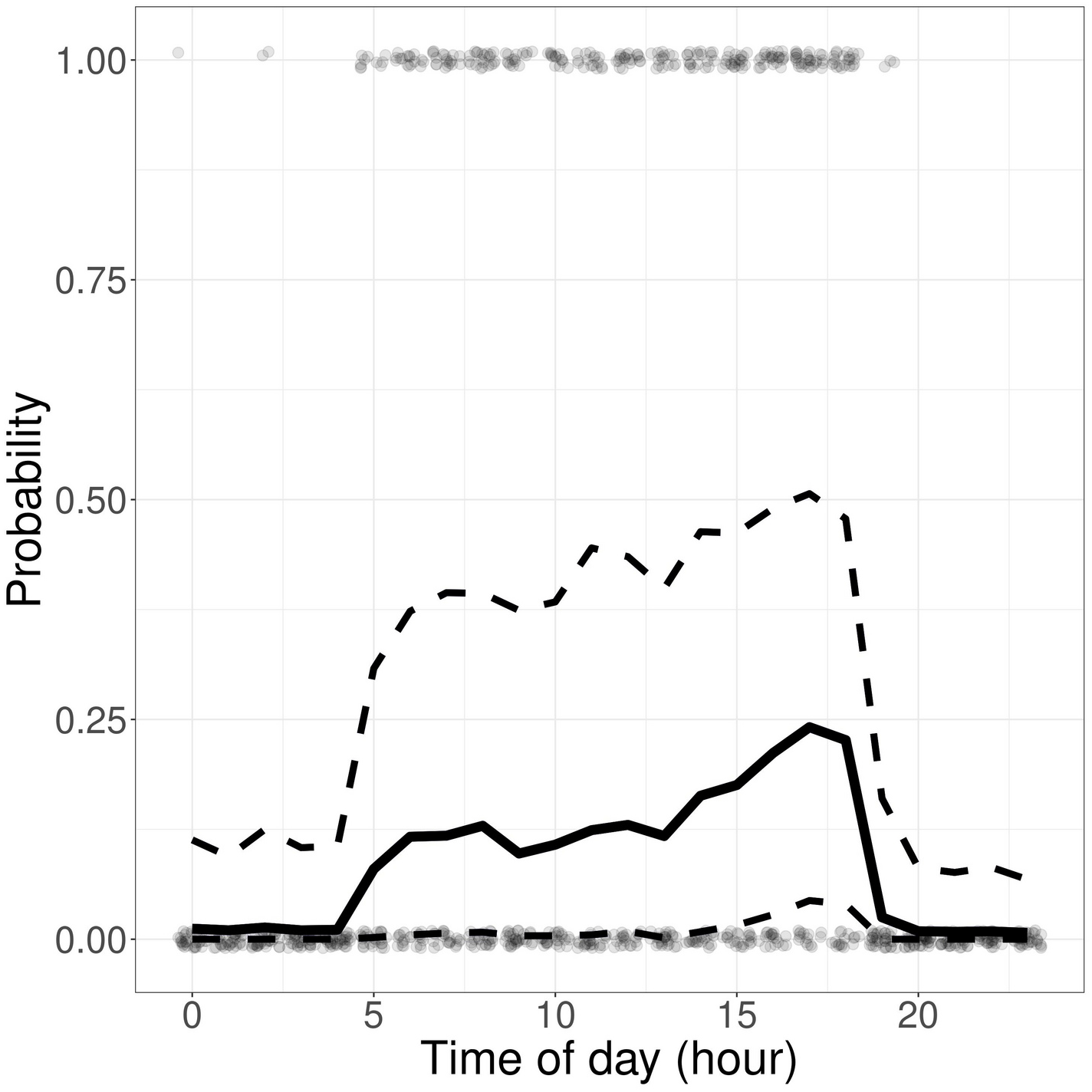
Probability of detecting a Saipan Reed Warbler vocalization by an experienced observer listening to 4 minutes of sound recordings randomly chosen from each hour. The solid line shows the model predictions from random forest models averaged over bootstrapped, cross-validated model replicates. Dashed lines show pointwise 95% bootstrap confidence intervals. Light grey dots show the observed detections (1) or non-detections (0) in 1 hour sound recordings. Light grey dots have been jittered horizontally and vertically to aid visualization. Saipan Reed Warblers sang during all daylight hours.

Time of day was a significant predictor of the probability of detecting a Saipan Reed Warbler vocalization; including time of day as a predictor variable significantly improved model prediction performance compared to a model with only nuisance covariates accounting for the identity of the observer and the sampling effort (model with time of day mean AUC = 0.735, 95% bootstrap CI [0.702, 0.764], mean Brier score = 0.184, 95% bootstrap CI [0.175, 0.194]; null model without time of day mean AUC = 0.508, 95% bootstrap CI [0.465, 0.553], mean Brier score = 0.227, 95% bootstrap CI [0.215, 0.239]). The mean predicted probability of detecting a Saipan Reed Warbler vocalization was highest in the afternoon around sunset, but confidence intervals around the predictions were wide (Fig. 1). The lower bound of the bootstrap 95% prediction interval remained close to zero at all times, except for the afternoon from about 1500 h until about 1900 h (Fig. 1).

## Discussion

Saipan Reed Warblers sang throughout the day, but rarely at night. There was no noticeable dawn chorus. Model results show some evidence that, if a Saipan Reed Warbler is present at a site, the probability of detection may be highest in the evening before sunset (Fig. 1, confidence interval not overlapping zero), but the wide confidence interval indicated that this pattern was weak.

Marshall et al. (2021) found no significant difference in Saipan Reed Warbler detection probability between in-person surveys conducted in the morning and in the afternoon, which is consistent with our findings. Frequent nighttime singing and a dawn chorus were reported in a short review summary (del Hoyo et al. 2020). We did not find any evidence for either of those phenomena, though we did detect a small number of vocalizations between midnight and sunrise (Fig. 1).

Surveying locations in both the morning and the afternoon could help maximize probability of detection. At all locations, Saipan Reed Warblers were detected in both morning and afternoon during this study. However, Saipan Reed Warblers were not always detected during both morning and afternoon on the same day or even during the same multi-day ARU deployment period; recordings from 2 locations had only afternoon detections during one of the deployment periods, and another location had only morning detections during one of the deployment periods. This suggests that, at some locations and some times, Saipan Reed Warblers might not vocalize all day long. It is possible that the variation in singing behaviors between locations and deployment periods is related to the breeding status of the birds. Flexible breeding schedules, with breeding possible all year long, is common in tropical birds (Stutchbury and Morton 2023). We did not attempt to determine the breeding status of birds in our study, so we do not have evidence linking singing behavior to breeding status. Saipan Reed Warblers have been detected nesting in all months except November and December (Mosher and Fancy 2002). Seasonal variation in territorial behavior and nesting has been reported for this species (Craig 1992, Mosher and Fancy 2002). Singing has been reported in all months (eBird Basic Dataset 2023). Our study design did not permit us to assess seasonal differences in singing behavior, but a study of seasonal singing activity would be useful for informing targeted surveys and for understanding the biology of this species.

The recommended USFWS survey protocols for Saipan Reed Warblers already include a possible survey window in the two hours before sunset, in addition to a morning survey window (USFWS 2014). However, our data suggest that surveys need not be limited to only morning and evening survey periods; observers targeting Saipan Reed Warblers may conduct surveys throughout the day. Additionally, we recommend that observers targeting Saipan Reed Warblers perform at least one site visit in the period before sunset, to maximize probability of detection if a Saipan Reed Warbler is present. Increasing the times of day during which surveys may be conducted gives more flexibility to government agencies and consultancies conducting surveys.

Surveys on Alamagan, which require logistically difficult and expensive expeditions, could also be conducted during any daylight hours, providing more flexibility and potentially greater sample sizes than would be possible if surveys are limited to mornings and before sunset. As an alternative to in-person surveys, automated recording units have the potential to cut down costly person-hours in the field. Extended duration ARU deployment on Alamagan, for example, could massively increase survey effort without incurring proportionally increasing costs associated with in-person expeditions to Alamagan. While automated detection algorithms would be a more efficient data processing tool, the method used in this study is an easily implementable, functional method for processing ARU recordings for Saipan Reed Warbler detections.

We noted some anecdotal differences between observers’ confidence when identifying Saipan Reed Warbler sounds. For example, some observers felt comfortable identifying call notes, while others did not. We did not attempt to distinguish calls from songs, but rather treated all vocalizations from Saipan Reed Warblers in the same way. Additional work will be useful to explore patterns in the types of vocal behavior, including calls, long-duration songs, and short song snippets. Of particular interest is the difference in vocal behavior between the sexes. Craig (1992) suggested that females do not sing. More recent observations have noted that females may call regularly (A.W. Santos, CNMI DFW, personal comm.). While we do not know of any well-documented cases of female song in Saipan Reed Warblers, female song is not unusual for sexually monomorphic tropical birds (Stutchbury and Morton 2023). It is possible that the lack of documented songs from females is due at least in part to the difficulty of visually determining the sex of Saipan Reed Warbler individuals, which have sexually monomorphic plumage. Genetic sexing, with accompanying behavioral observations, could helpfully distinguish male and female vocal behavior, which would improve our understanding of the differences in vocal behavior and detectability between sexes.

Singing activity in Saipan Reed Warblers was strongly limited to daylight hours in our study, but this does not necessarily imply that the birds are not active at night. Mukhin et al. (2009) found that Common Reed Warblers (*Acrocephalus scirpaceous*) made nocturnal flights in response to simulated nest loss, and also when experimentally moved away from breeding territories during the breeding season. We are not aware of any studies tracking nighttime movements of Saipan Reed Warblers, so the possibility remains that the birds are moving around but not vocalizing during the night. For the purposes of targeted surveys, most detections of Saipan Reed Warblers are by sound, so focusing survey effort during daylight hours when the birds are vocalizing makes sense.

## Acknowledgments

ADA Tudela assisted with data collection. We thank AW Santos for informative conversations about Saipan Reed Warblers, and RJ Craig for useful comments on an earlier draft of this manuscript. This work was supported by the U.S. Fish and Wildlife Service through the Wildlife and Sport Fish Restoration Grant Program, under grant number F22AF03625. This project was supported by grants from the National Science Foundation (NSF), Louis Stokes Alliances for Minority Participation (LSAMP), Award number: 1826864. The content is solely the responsibility of the authors and does not necessarily represent the official views of the National Science Foundation.

## Notes

https://doi.org/10.5281/zenodo.10586984

### Competing Interest Statement

The authors have declared no competing interest.

