## Supplementary material for "Daily vocalization patterns of the Saipan Reed Warbler (*Acrocephalus hiwae*)": Supplelmental Table S1

Supplemental Table S1: Number of surveys for Saipan Reed Warbler conducted by the CNMI Division of Fish and Wildlife at proposed development sites during each hour of the day, from 2005-2023.

| **Hour of Day** | **Number of Surveys** |
|---|---|
| 0 | 0 |
| 1 | 0 |
| 2 | 0 |
| 3 | 0 |
| 4 | 3 |
| 5 | 6 |
| 6 | 99 |
| 7 | 814 |
| 8 | 1384 |
| 9 | 1384 |
| 10 | 773 |
| 11 | 157 |
| 12 | 60 |
| 13 | 428 |
| 14 | 464 |
| 15 | 196 |
| 16 | 14 |
| 17 | 0 |
| 18 | 0 |
| 19 | 0 |
| 20 | 0 |
| 21 | 0 |
| 22 | 1 |
| 23 | 0 |
